# Biological hydrogen cyanide emission globally impacts the physiology of both HCN-emitting and HCN-perceiving *Pseudomonas*

**DOI:** 10.1101/2021.09.29.462390

**Authors:** Abhishek Anand, Laurent Falquet, Eliane Abou-Mansour, Floriane L’Haridon, Christoph Keel, Laure Weisskopf

## Abstract

Bacterial volatile compounds have emerged as important chemical messengers between bacteria themselves as well as in their interactions with other organisms. One of the earliest examples of bioactive volatiles emitted by bacteria is hydrogen cyanide (HCN), which was long considered a mere respiratory toxin conferring competitive advantage to cyanogenic strains. Using cyanide-deficient mutants in two *Pseudomonas* strains and global transcriptome analysis, we demonstrate that the impact of HCN is much more global than previously thought. We first observed that the lack of cyanogenesis in emitting strains led to massive transcriptome reprogramming affecting diverse traits such as motility and biofilm formation (respectively inhibited vs. promoted by HCN), or the production of siderophores, phenazines and other antimicrobial compounds (repressed by HCN). We then exposed non-cyanogenic strains to biogenically emitted HCN from neighboring cells and observed similar transcriptome modulations and phenotypic changes, suggesting that HCN not only acts endogenously but also exogenously, remotely manipulating important traits involved in competition and virulence, e.g. siderophore production, in other organisms. Cyanogenesis in *Pseudomonas* has long been known to play a role in both the virulence of opportunistic pathogens and the efficient biocontrol activity of plant-beneficial strains, however this impact was so far thought to occur solely through the inhibition of respiration. We demonstrate here new ecological roles for a small and fast-diffusing volatile compound, which opens novel avenues in our understanding of and ability to interfere with important processes taking place in pathogenic and beneficial *Pseudomonas* strains.

**Importance:** Bacteria communicate by exchanging chemical signals, some of which are volatile and can remotely reach other organisms. Hydrogen cyanide (HCN) was one of the first volatiles discovered to severely impact exposed organisms by inhibiting their respiration. Using HCN-deficient mutants in two *Pseudomonas* strains, we demonstrate that HCN’s impact goes beyond the sole inhibition of respiration and affects both emitting and receiving bacteria in a global way, modulating their motility, biofilm formation and production of antimicrobial compounds. Our data suggest that bacteria could use HCN not only to control their own cellular functions, but also to remotely influence the behavior of other bacteria sharing the same environment. Since HCN emission occurs in both clinically and environmentally relevant *Pseudomonas*, these findings are important to better understand or even modulate the expression of bacterial traits involved in both virulence of opportunistic pathogens and in biocontrol efficacy of plant-beneficial strains.

## Introduction

The exchange of volatile compounds as means of communication between bacteria and other organisms has recently gained increasing interest over the last two decades (1). One of the first volatiles discovered to play a role in biotic interactions is hydrogen cyanide (HCN), a well-known toxin irreversibly binding to the key respiratory enzyme cytochrome C oxidase. Cyanogenesis occurs in many different types of organisms, including bacteria, plants, animals and even humans (2–4). In bacteria, cyanogenesis occurs in different but restricted taxa such as *Proteobacteria* (e.g. *Chromobacterium, Pseudomonas, Rhizobium*) or cyanobacteria (e.g. *Anacystis, Nostoc*), although most studies have focused on *Pseudomonas* strains, which form hydrogen cyanide (HCN) by oxidative decarboxylation of glycine (5). Cyanogenic strains have evolved a range of mechanisms which enable them to avoid the toxic effects of their own product, for instance the enzymatic conversion of HCN to the less harmful thiocyanate (6), or the reliance on alternative cyanide-insensitive cytochrome c oxidases (7–9). HCN is considered an important secondary metabolite in both environmental (e.g. *Pseudomonas protegens* CHA0) and clinical (e.g. *Pseudomonas aeruginosa* PAO1) strains (10, 11). In the clinical context, HCN emission has been shown to occur in the lungs of cystic fibrosis patients harbouring *P. aeruginosa* strains and to significantly affect lung function (12). In addition to this direct effect on the host, cyanogenesis was recently shown to confer a competitive advantage to lung-infecting *P. aeruginosa* via the inhibition of co-infecting *Staphylococcus aureus*, a frequent co-inhabitant of cystic fibrosis patient lungs (13).

In the environmental context too, the ecological role of HCN emission in bacterial defense against competing organisms is well established and this molecule has been repeatedly pinpointed as a major actor in the arsenal of root-associated *Pseudomonas* with protective effects against plant pathogens (14). In an earlier study using comparative genomics to identify the molecular determinants of late blight disease control by potato-associated *Pseudomonas*, HCN came out as a likely candidate accounting for the inhibition of the devastating oomycete pathogen *Phytophthora infestans* (15). However, when comparing the anti-*Phytophthora* effects of cyanogenesis-deficient mutants in two different species of *Pseudomonas* with that of their respective wild types, only little to no reduction in their antagonistic potential was observed. On the contrary, the mutants seemed to have even gained additional antimicrobial capacities, since they induced higher inhibition of the pathogen’s zoospore germination compared with the wild types (16). This observation raised the question wether HCN, in addition to acting on neighboring organisms, was also influencing the physiology of the emitting bacteria themselves, as previously shown for other so-called “secondary metabolites” (e.g. 2,4-diacetylphloroglucinol (17) or phenazines (18)) initially discovered for their impact on target organisms but later demonstrated to fulfill important physiological functions in the producing bacteria. In order to answer this question, we took advantage of previously generated HCN-deficient mutants in two different strains of plant-associated *Pseudomonas* and first compared their transcriptome to that of their repective wild types. We then investigated whether the changes observed were restricted to endogenous effects on the emitting bacteria themselves, or whether they would also be triggered remotely in neighboring bacteria exposed to biogenically produced HCN.

## Results and discussion

In a former study quantifying the impact of HCN emission on the biocontrol activity of potato-associated *Pseudomonas* strains against *Phytophthora infestans* zoospore germination, we made the unexpected observation that the HCN-deficient mutants displayed stronger inhibitory activities compared with their respective wild types (16). As iron deprivation mediated by the excretion of chelating agents is a well-known mechanism of antifungal activity (19), we started the present work by quantifying the mutants’ production of siderophores. We first grew both strains on King’s B medium, which is classically used to detect siderophores in pseudomonads. In contrast to the negative control *Escherichia coli*, our two model strains *Pseudomonas chlororaphis* R47 and *Pseudomonas putida* R32 secreted siderophores as evidenced by the halo surrounding the colonies (Figure 1a). In *Pseudomonas* R32, the cyanide-deficient mutant indeed appeared to produce more siderophores than the wild type, while the difference was less clear for R47 in this qualitative plate assay. Since we knew from these two strains’ genomes that they both encode pyoverdine production, we used a fluorospectrometry-based assay (20) to quantify the production dynamics of this particular siderophore as well as the growth of both wild types and mutants in liquid King’s B medium (Figure 1b). The production of pyoverdine was significantly higher in both cyanide-deficient mutants compared with the corresponding wild type strains. For R47, faster growth was observed for the mutant at the onset of the stationary phase, yet the strong increase in pyoverdine production was in no proportion to this moderate growth advantage (Figure 1b). For R32, the decreased growth of the wild type compared with the mutant was less striking. This differential impact of cyanogenesis on growth in the two *Pseudomonas* strains might be linked to their respective HCN production levels, which were higher (ca. 80 µM) for R47 than for R32 (ca. 28 µM) (Figure S1). The orange color of R47 (Figure 1a) originates from the production of phenazines, an important class of antimicrobial compounds, also involved in iron acquisition, redox balance, and many other biological functions in *Pseudomonas* strains (21). We therefore also quantified the production of phenazine carboxylic acid (PCA), one particular form of phenazines, in both wild type and cyanide-deficient mutant strains grown in King’s B medium. We analyzed this phenotype only in *P. chlororaphis* R47, since phenazine biosynthetic genes are absent from the genome of *P. putida* R32. We observed that similarly to pyoverdine, PCA was also produced at a higher concentration (>2.5-fold) by the mutant than by the wild type (Figure 1c). Next to PCA, other phenazines were also produced more abundantly in the cyanide-deficient mutant (Figure S2). Taken together, these data indicate that the absence of HCN led to a significantly enhanced production of other secondary metabolites that could fulfil similar ecological functions, such as the inhibition of microbial competitors.

**Figure 1.**
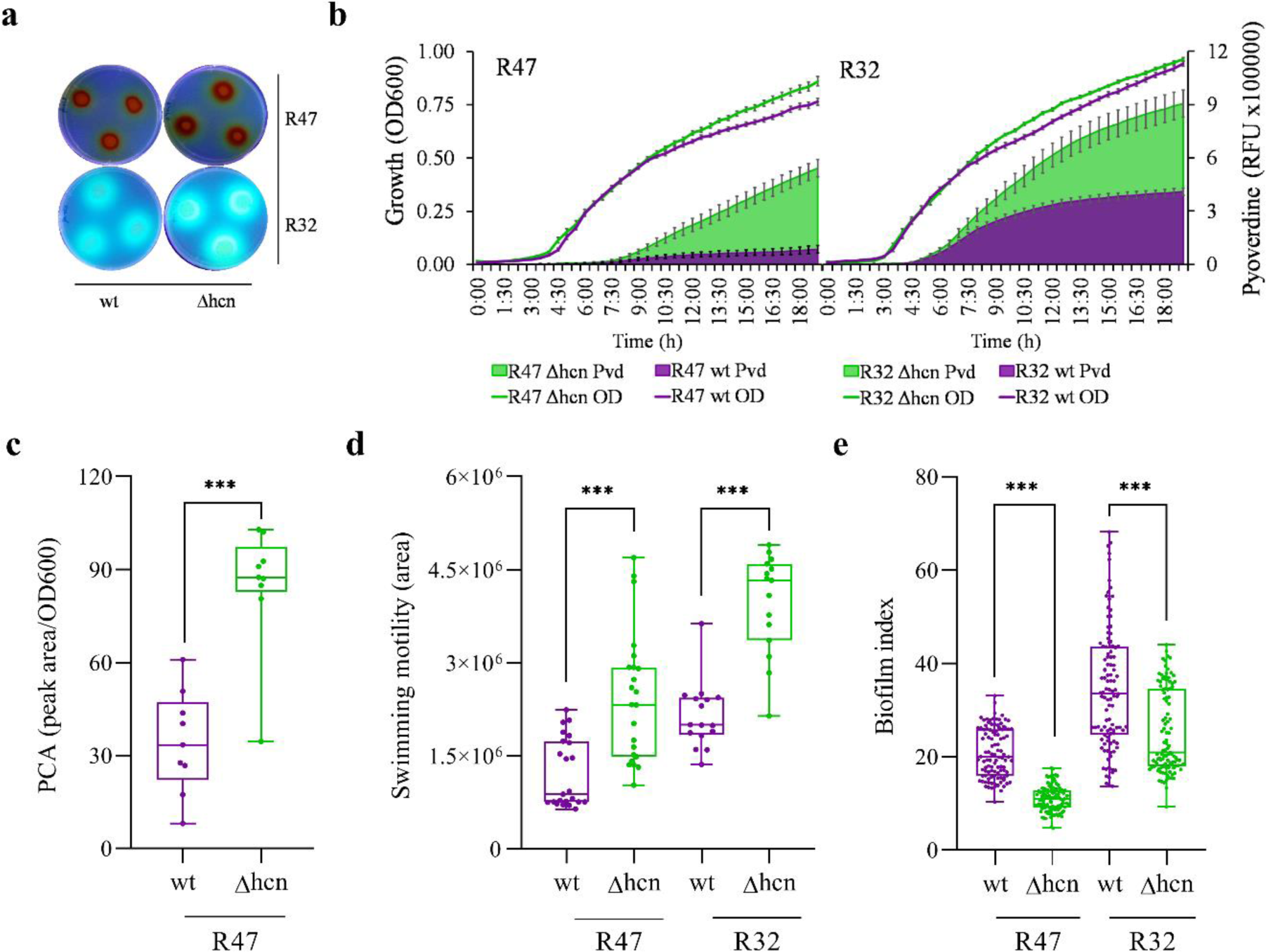
Loss of cyanogenesis increases pyoverdine and phenazine production, induces swimming motility and represses biofilm production. a, Qualitative analysis of siderophore production by wild type (wt) and cyanide-deficient (Δhcn) mutant strains of both *Pseudomonas chlororaphis* R47 and *Pseudomonas putida* R32 after five days of growth at room temperature on King’s B agar medium. Plates were visualized under UV light. b, Growth curves (optical density (OD) at 600 nm, lines) and pyoverdine (Pvd) detection assay (fluorescence reading at 465nm, bars) in liquid King’s B medium grown for 19h at 30 °C. Data represent the average of three independent experiments with four technical replicates each. c, Phenazine quantification assay based on HPLC-UV detection. R47 wt and R47 Δhcn strains were grown at 30 °C in liquid King’s B medium for 24h before sampling. Bars represent the phenazine carboxylic acid (PCA) peak area normalized with growth (OD600). Data represent the average of three independent experiments with three technical replicates each. d, Swimming motility assay on low agar M9 plates. The colony area was measured using the ImageJ software after four to five days incubation at room temperature. Bars show the average of four and three independent experiments for R47 and R32, respectively, with four to six technical replicates each. e, Biofilm formation assay. Strains were grown statically in liquid King’s B medium in 96-well plates for 48h at 30 °C. Bars correspond to biofilm index (crystal violet staining measured at 570 nm divided by optical density measured at 550 nm before the staining) and represent the average of three independent experiments with 36 technical replicates each. For c-e, samples were statistically analyzed using Mann-Whitney U test (two-tailed). ***P<0.001. Error bars represent standard error.

The ability to move towards a favorable environment and once there, to form a biofilm are relevant features for successful colonization of both plant (e.g. root) and human (e.g. lung) tissues by pseudomonads (22, 23). We therefore wondered whether motility and biofilm formation would also be influenced by the loss of cyanogenesis in the two *Pseudomonas* mutants. Swimming motility assays revealed that both cyanide-deficient mutants were significantly more motile (ca. 2-fold) than the wild types (Figure 1d). In contrast to all previous phenotypes tested, where the mutants performed better than the wild types, biofilm formation was significantly lower for both mutants compared to the wild types (Figure 1e). In addition to confirming the expected negative relation between a motile and a sessile life style, these data provided the proof of concept that HCN could also stimulate and not only inhibit physiological processes in cyanogenic pseudomonads, suggesting that its effects may go beyond those of a mere toxin. Overall, the consistent observation in two different *Pseudomonas* species that the loss of cyanogenesis led to effects as diverse as enhanced siderophore production and reduced biofilm formation, confirmed our hypothesis that HCN likely has a global impact on bacterial physiology and behavior.

In order to test this hypothesis, we selected *P. chlororaphis* R47 because of its large genome encoding many secondary metabolites (15) and compared the transcriptome of the wild type and that of the cyanide-deficient mutant grown in liquid King’s B medium at three different time points (Figure 2a). We chose these time points to cover i) the early log phase, at which we did not expect any change since HCN, as a secondary metabolite, was classically reported to be produced at the end of the exponential growth phase (24), ii) the transition between exponential and stationary growth phase, at which HCN effects were expected to start taking place but no difference in growth was yet observed between the wild type and the mutant, and iii) the later stationary growth phase, at which we expected changes to be further amplified but where they would likely be confounded with different growth dynamics (Figure 2a). Contrary to our expectations, the transcriptomes of the wild type and the cyanide-deficient mutant clustered separately already at the early log phase (Figure 2b). Looking at the expression of the HCN biosynthetic operon in *P. chlororaphis* R47 wild type, we observed that it was indeed expressed at the early exponential phase already (Figure S3), and its absence induced the dysregulation of 32 genes belonging to different COG categories at this early time point (Figure 2c and Table S1). Among these, genes involved in signal transduction and lipid transport / metabolism were mainly upregulated in the absence of HCN, while genes involved in transcription, amino acid and carbohydrate metabolism, inorganic ion transport and secondary metabolism were mainly downregulated. Genes belonging to the energy production and conversion category were also dysregulated, with some being up- and others downregulated in the mutant, such as the genes encoding cytochrome c oxidase orphan subunits. The early expression of alternative oxidases in the wild type compared with the mutant constitutes a further proof that cyanogenesis occurs earlier than classically assumed and is not solely conditioned by the lack of nutrients or of oxygen occurring at the early stationary phase, where HCN production has been previously reported as being maximal (5).

**Figure 2.**
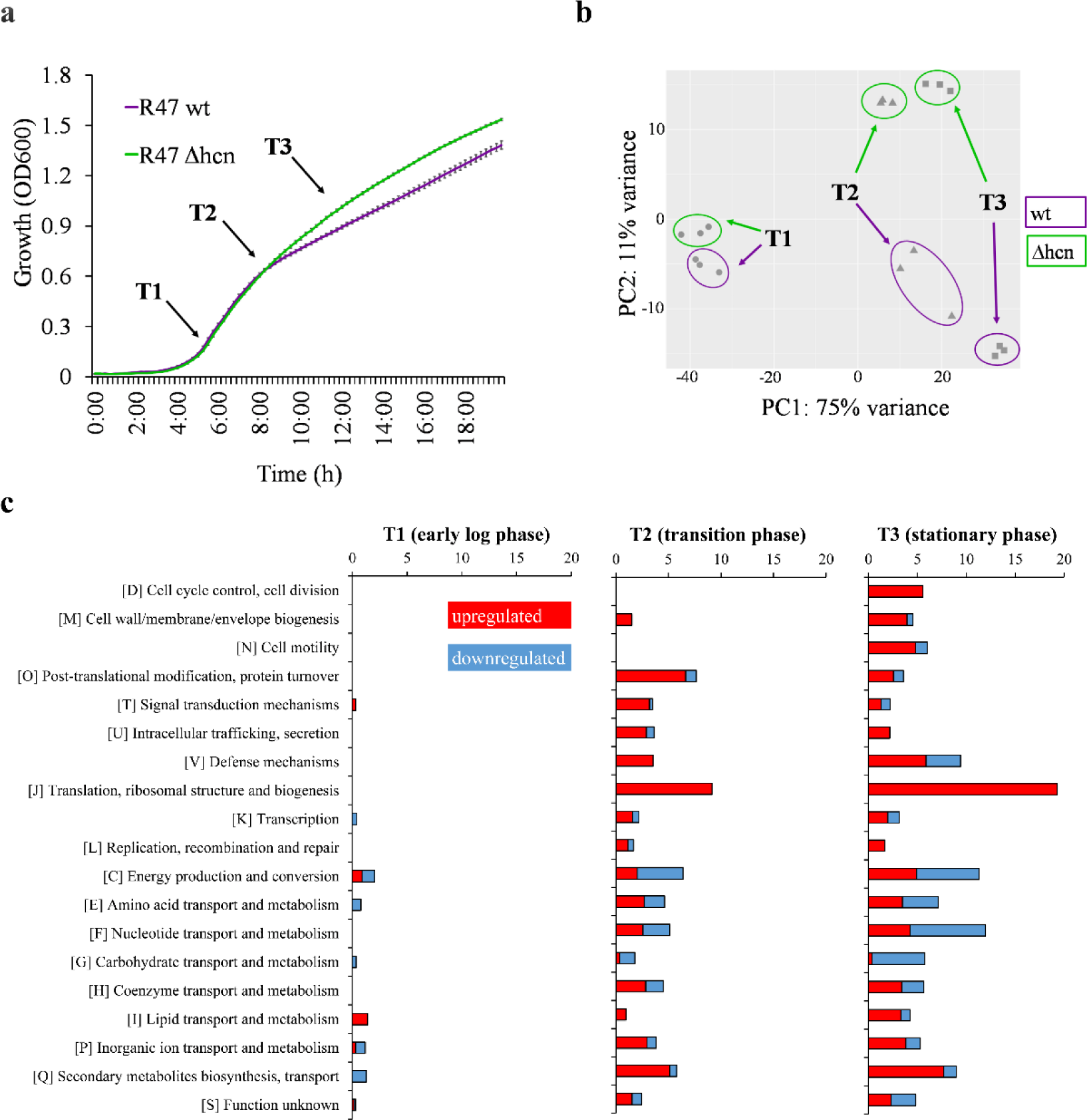
Inability to produce HCN leads to global transcriptional dysregulation. a, Representative growth curve of *Pseudomonas chlororaphis* R47 wt and Δhcn strains in King’s B liquid medium highlighting the three time points used for transcriptome analysis. T1; early log phase, T2; transition phase (exponential to stationary), T3; stationary phase. b, Principal component analysis of transcriptomic data at the three time points. Strains were grown in King’s B liquid medium in 6-well plates at 30 °C with continuous shaking before each sampling. c, COG category analysis of the dysregulated genes in R47 Δhcn strain compared with its wild type. Bars show the percentage of dysregulated genes in each COG category. RNA-sequencing data (b,c) were obtained from three independent experiments. Cut-off for significance; Log2 (fold change) ≥ 1, q-value ≤ 0.05.

As expected, differences between the transcriptomes of the wild type and the mutant increased over time (Figure 2b,c), with 206 genes (ca. 3.2 % of the genome) dysregulated at the transition between the exponential and stationary growth phases, and 350 (ca. 5.5 %) at the late stationary growth phase, where all COG categories were affected by the presence vs. absence of HCN. These genome-wide expression changes already started at the second time point (transition to stationary growth phase), where both strains still followed the same growth dynamic (Figure 2a), which suggests that the wild type was not or only marginally affected by HCN toxicity at this growth stage but still showed genome-wide transcription changes compared with the cyanide-deficient mutant. Overall, the absence of HCN in the mutant led to upregulation of many genes, as for instance those involved in pyoverdine and phenazine synthesis (Figure 2b,c and Table S2), which corroborated our phenotypic assays (Figure 1b,c and Figure S2). Among the few genes upregulated by HCN, one was predicted to encode the helix-turn-helix-type transcriptional regulator MlrA which was strongly downregulated in the cyanide-deficient mutant (Table S2). In *E. coli* and *S. enterica*, MlrA was shown to activate extracellular matrix production (25). This could explain the higher biofilm forming capacity observed in the *Pseudomonas* wild types compared to the cyanide-deficient mutants in the present study (Figure 1e). Higher biofilm formation by wild types could also be explained by higher expression of the type VI secretion system (T6SS) (Table S2). This system has been shown to be involved in biofilm formation by *P. aeruginosa* (26–28). Next to its putative role in biofilm formation, the T6SS has been involved in bacterial interactions with competitors (29), similarly to HCN itself. Likewise, the operon encoding the insecticidal Fit toxin (30) was also upregulated in the absence of HCN (Table S2), suggesting a possible compensation of the lack of one biological weapon with the higher production of others (e.g. phenazines, pyoverdine, Fit toxin and T6SS). Regarding motility, transcriptomic analysis revealed upregulation of flagellar biosynthesis genes in the cyanide-deficient mutant (Table S2), which could explain both the mutant’s higher motility and lower biofilm formation (Figure 1d,e). Altogether, the global transcriptome shifts caused by the presence vs. absence of HCN corroborated the phenotypic differences observed between the wild type and the cyanide-deficient mutants (Figure 1) and highlighted endogenous HCN as a global regulator of gene expression, compared e.g. with the global activator of secondary metabolism (GacA), whose mutation led to ca 13 % of dysregulated genes in the closely related strain *P. chlororaphis* 30-84 (31). However, the extent of the observed HCN-induced transcriptome changes is in stark contrast to earlier work on *P. aeruginosa*, which reported only 16 dysregulated genes in an cyanide-deficient mutant (9). This discrepancy might be due to a higher resolution of RNA-Seq over microarray analysis, or to different culture conditions, with the complex but iron-limited King’s B medium in our case and glycine-enriched minimal medium boosting HCN emission in the earlier study. Interestingly, many of the phenotypes we observed in the mutants can be linked to iron availability: siderophore and phenazine production are evident examples, but also motility and biofilm formation have been linked to iron availability in *Pseudomonas aeruginosa* (32). While iron was shown to inhibit motility and to increase biofilm formation in this bacterium, we observed a similar effect of HCN itself in our experiments (with decreased motility and increased biofilm in the wild type), although HCN is rather expected to bind iron and therefore to reduce its availability (33). The higher emission of siderophores in the HCN mutant is also consistent with the hypothesis that endogenously produced HCN would not act by scavenging iron, which would result in the wild type needing higher siderophore production than the cyanide-deficient mutant. This does not rule out an interaction between iron and HCN, especially since iron itself was shown to increase HCN emission in different cyanogenic bacteria (5). This direct effect of iron on cyanogenesis makes it difficult to disentangle iron- and HCN-mediated effects, as growing the strains under different iron regimes would lead to differential HCN emission.

Since HCN is a gaseous molecule easily diffusing into the vicinity of cyanogenic strains (Figure S4), we wondered whether neighboring, non-cyanogenic cells would respond to exogenous HCN supply with similar changes in physiology as those we observed between mutants and wild types. We therefore carried out an experiment where the cyanide-deficient mutants of R47 and R32 were grown in volatile-mediated contact with their respective wild type (Figure 3a), and compared the phenotypes of the wild types, the mutants and the HCN-exposed mutants. To exclude a volatile-mediated effect that would not be caused by HCN itself but by other volatile compounds exhibiting altered emission in the mutant compared with the wild type, we first verified that the volatile profiles of both strains exhibited no other significant difference except for the presence vs. absence of HCN (Figure S5). Exposing the mutants of both R47 and R32 to exogenous HCN emitted by their respective wild types did not impair the mutants’ growth (as would be expected if HCN would be toxic at these concentrations), yet it suppressed the increase in pyoverdine production observed in the non-exposed mutants (Figure 3b) and restored both mutants’ biofilm formation and swimming motility to the wild type levels (Figure 3c,d). For the motility rescue experiment, in the R32 cyanide-deficient mutant, we observed swarming rather than swimming motility in the chosen split plate setup (Figure 3d). We then also compared the transcriptomes of exposed and non-exposed mutants of R47 with that of the wild type to see whether the phenotypes induced by exogenous exposure to biologically produced HCN would be corroborated by changes at the gene expression level (Figure 3e). At the early exponential phase, both non-exposed and exposed mutants of R47 showed similar transcriptomes (only four differentially expressed genes, see Figure S6 and Table S1), suggesting that at this early time point, HCN was affecting only the producing cells but was not yet reaching relevant concentrations in the neighboring wells. However, at the later time point corresponding to the transition between the exponential and stationary growth phases, most of the gene expression changes observed in the mutant were rescued by exposure to exogenous HCN (Figure 3e, Figure S6b,c). These included the changes in genes encoding pyoverdine and phenazine synthesis, the insecticidal Fit toxin, as well as the MlrA activator, the T6SS and the flagellar biosynthetic genes potentially involved in biofilm formation, which corroborated the rescued phenotypes (Figure 3b,c,d and Table S2). The transcriptome changes observed at this intermediate time point between the mutant and the exposed mutant mimicked those observed between the mutant and the wild type, confirming that the exposed mutant behaved largely like a wild type (Figure 3b,e). Principal component analysis showed that this transcriptome similarity was further increased at the late stationary phase (time point 3), although the number of dysregulated genes increased from 19 to 141, which might be due to the differential growth between the wild type and the exposed mutant observed at this later time point (Figure 3b, Figure S6a,b). Interestingly, the proportion of genes rescued by external supply of HCN at the transition between the exponential and stationary growth phases differed between the COG categories, with a lower rescue proportion in the categories involved in housekeeping functions and primary metabolism (e.g. ribosome structure, replication, energy production, or carbohydrate, amino acid and nucleotide metabolism) than in those potentially involved in biotic interactions (e.g. defense mechanisms, inorganic ion transport and secondary metabolism) (Figure S6c).

**Figure 3.**
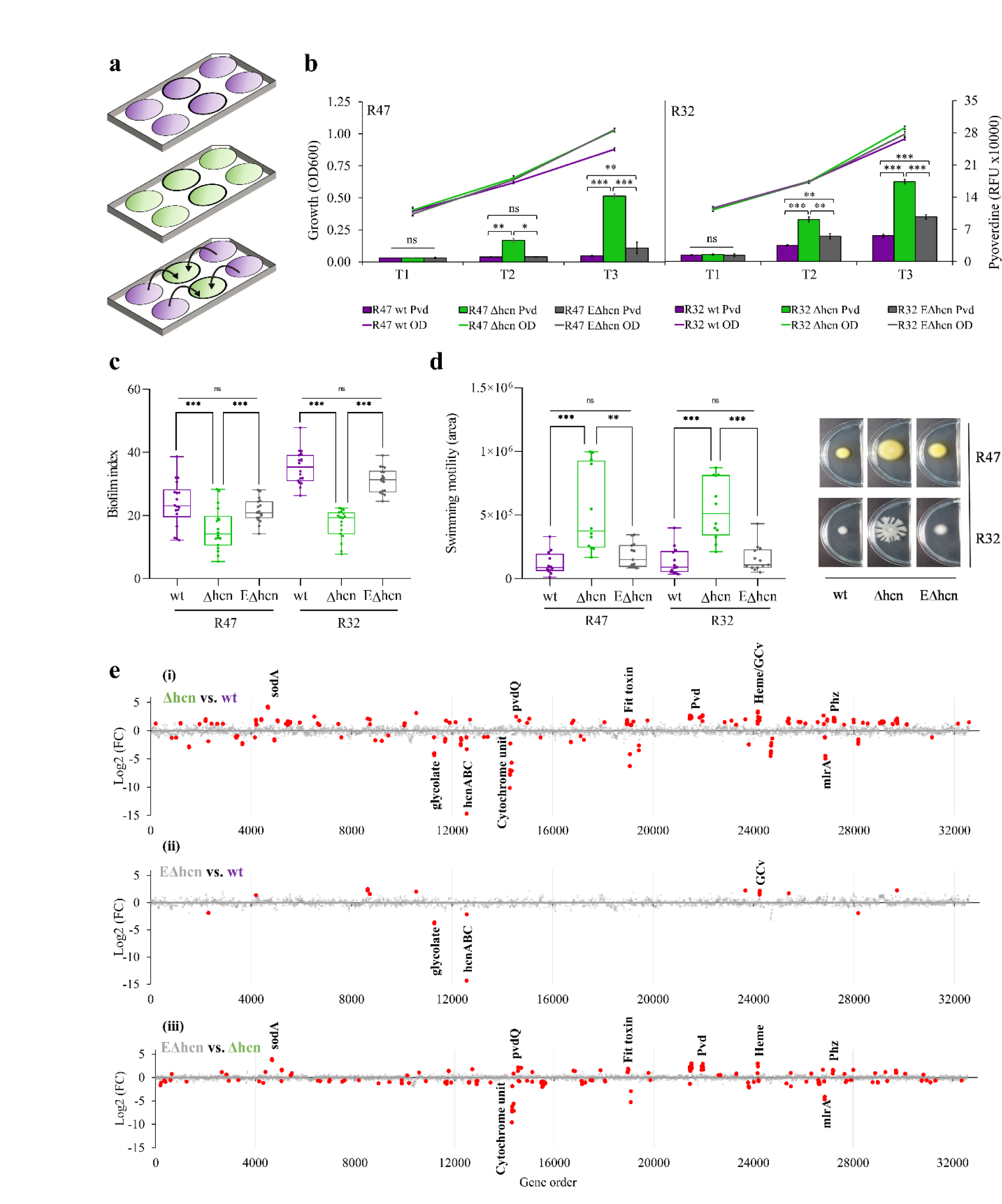
Exposure to exogenous HCN emitted by *Pseudomonas chlororaphis* R47 and *Pseudomonas putida* R32 wild types represses pyoverdine production and swimming motility, but induces biofilm formation in the cyanide-deficient mutants. a, Scheme of the volatile-mediated exposure experiment using 6-well plates used for (b). Purple color represents liquid cultures of the wild type, green color liquid cultures of the cyanide-deficient mutants. The middle two wells (thicker border) were collected for RNA extraction and sequencing. b, Growth (OD600) and pyoverdine (pvd) measurements for *P. chlororaphis* R47 and *Pseudomonas putida* R32 strains grown at 30 °C in liquid King’s B medium in 6-well plates. wt, wildtype; Δhcn, cyanide-deficient mutant; EΔhcn, Δhcn strains exposed to HCN emitted by the respective wild types. Measurements were performed as indicated in Figure 1. Data represent the average of three independent experiments with two technical replicates each. T1 (early log phase), T2 (transition phase between exponential and stationary phases) and T3 (stationary phase). c, Biofilm formation assay for wild types (wt), cyanide-deficient mutants (Δhcn) and HCN-exposed mutants (EΔhcn). The assay was carried out in liquid LB as indicated in Figure 1 and Figure S8. Bars represent the average of three independent experiments with 6-20 technical replicates each. d, Swimming motility assay on split Petri dishes. The wild type and the cyanide-deficient mutant were grown on each side of the split Petri dishes (Figure S7). Plates were incubated at room temperature for two days before pictures were taken. The areas were measured as indicated in Figure 1. Bars show averages of three independent experiments with four to five technical replicates each. e, Genome-wide representation of differentially expressed genes at timepoint 2 (transition phase between exponential and stationary growth phase). The x-axis shows the gene location on the chromosome with each dot corresponding to one gene and the y-axis shows the log2 (fold change) values. (i) and (ii) log2 (fold change) values correspond to Δhcn and EΔhcn, using the wt as reference. (iii) Log2 (fold change) values correspond to the Δhcn strain using EΔhcn as reference. Red dots represent significantly differentially expressed genes while grey dots represent no significant difference as compared to the respective reference. Cut-off for significance; Log2 (fold change) ≥ 1, q-value ≤ 0.05. For b-d, samples were statistically analyzed using ANOVA followed by Tukey’s multiple comparison test. Error bars represent the standard error. *\*\*\*P*<0.001, *\*\*P*<0.01, *\*P*<0.05; ns, not significant. For e, dots represent average values from three independent experiments for each time point and treatment. SodA, superoxide dismutase A; hcnABC, hydrogen cyanide synthase ABC; cytochrome unit, cytochrome orphan subunit N4Q4; Pvd, pyoverdine biosynthesis; Phz, phenazine biosynthesis; glycolate, glycolate to glyoxylate oxidation; heme, heme utilization; mlrA, activator of csgD (master regulator of biofilm formation); GCv, glycine-cleavage system.

Altogether, our data indicate that the biological functions of HCN are much wider than previously thought. Similarly to phenazines, which were also first categorized as “mere secondary metabolites” and later turned out to act as internal signaling molecules with diverse and important cellular functions (34), we provide evidence here that HCN is much more than a respiratory toxin, and acts as an inter- and extracellular signal on both producing and receiving cells. These findings are in line with recent reports on the ability of HCN to induce broad physiological effects in both plant and human cells (35, 36). Compared with phenazines, HCN is a volatile molecule and as such, its zone of influence is expected to be much broader (1). Our data show that this small volatile compound remotely leads to global changes in the behavior of neighboring bacteria, repressing their iron uptake, swimming motility and synthesis of biological weapons such as phenazines, the insecticidal Fit toxin or the type VI secretion system, which likely confers a substantial advantage to cyanogenic bacteria over non-cyanogenic ones in the highly competitive environments they usually colonize, from the plant rhizosphere to the human lung.

## Experimental procedures

### Strains and culture media

*Pseudomonas chlororaphis* (R47) and *Pseudomonas putida* (R32) and their derived Δhcn mutants were grown at 30 °C on lysogeny broth (LB) agar medium (20 g/L of LB mix (Lennox, Fisher Bioreagents) with 15 g/L of agar (Agar-agar, Kobe I, Carl Roth)) (37). The generation of Δhcn strains has been previously described (16). King’s B (KB) medium was prepared by mixing 20 g/L of proteose peptone #3 (Bacto^TM^ Proteose Peptone No. 3, ThermoFisher Scientific), 1.5 g/L of dipotassium phosphate (Carl Roth GmbH + Co. KG), 1.5 g/L of magnesium sulfate heptahydrate (Carl Roth GmbH + Co. KG) and 10 ml/L of glycerol (Carl Roth GmbH + Co. KG) and 15 g/L of agar (when needed).

### Growth curves and pyoverdine measurements

Bacterial colonies (two to three single colonies) were used to inoculate LB and cultured for 19h at 30 °C with shaking (180 rpm). The cells were collected by centrifugation at 5000 rpm for five min and then washed twice with 0.9 % NaCl. The bacterial suspension was then adjusted to OD600 = 1 (henceforth named ‘OD adjusted’) and 5 µl was used to inoculate 100 µl of KB broth in 96-well plates (Corning, NY, USA). Parafilm was used to seal the plates. A Cytation5 automated plate reader (BioTek, Winooski, VT, USA) was used to incubate plates at 30 °C with continuous shaking. Reads were taken at 600 nm to generate growth curves and pyoverdine measurements were done at excitation = 405 nm and emission = 460 nm (20). For HCN exposure experiments, similar readings were taken using 6-well plates (Corning, NY, USA) (Figure 3a). Three ml KB broth were added to each well and 300 µl of OD adjusted bacterial suspension were added. Plates were incubated at 30 °C with shaking at 90 rpm and readings were taken for 5, 7 and 11h. All experiments were replicated three times independently in total.

### Phenazine measurement and identification

Phenazines were quantified as previously described using a high-performance liquid chromatography (HPLC) method (38). Briefly, OD adjusted bacterial cell suspensions were used to inoculate 3 ml of KB broth and incubated at 30 °C with shaking at 180 rpm for 24h. The cultures were then filtered using 0.2-µm filters (Fisher Scientific International, Inc.) and stored at −20 °C until analysis. Analyses were performed by a HPLC equipped with a quaternary pump P100A, coupled with a UV diode array detector ultimate 3000 (Dionex, Thermoscientific, Olten, Switzerland). The samples (40 μl) were injected onto a MN Nucleosil C18 analytical column (250 mm x 4 mm i.d., 100-5) (Macherey-Nagel, Duren, Germany), using a flow rate of 0.8 mL.min^-1^ at 27 °C. The mobile phase consisted of solvent A (0.1% TFA (trifluoroacetic acid) in water) and solvent B (0.1 % TFA in acetonitrile); the gradient started with 5 % of solvent B and reached 15 % at 2 min, 83 % at 15 min, 95 % at 16 min, and 100 % at 45 min. Phenazines were detected at 365 and 250 nm with diode array on-line detection. UV spectra were recorded between 200 and 800 nm. For compound identification, samples were freeze dried, dissolved in 500 μl of ethyl acetate and purified by preparative TLC on silica gel (Kieselgel 60, F_254_, 0.25, Merck, Darmstadt, Germany) using CHCl_3_:MeOH (95:5) as the eluent. The spots were visualized by exposure to UV radiation at 256 nm. Three compounds were isolated and identified by HPLC and mass spectrometry as 1 or 2-hydroxyphenazine (i) (Rf 0.72; RT 14.5; m/z 197.07 [M+H]^+^), phenazine-1-carboxylic acid (PCA) (ii) (Rf 0.64; RT 16.4; m/z 225.06 [M+H]^+^) and 2-hydroxyphenazine carboxylic acid (iii) (Rf 0.58; RT 21.6; m/z 241.06 [M+H]^+^). ESI-MS/MS analysis in positive ionization mode was performed on Q Exactive Plus mass spectrometer (Thermo Fisher Scientific). Mass spectra were recorded between 50 and 500 *m/z*. The peak areas at the specific wavelength l_max_ 250 nm were chosen for qualitative quantification. Three biologically independent replicates were collected with three technical replicates in each experiment.

### Swimming motility assays

Swimming motility assays were performed as previously reported (39). Briefly, OD adjusted inoculum was pricked into 0.3 % M9 agar medium (For 1 liter, 10 ml of 20 % glucose (Carl Roth GmbH + Co. KG), 10 ml of 20 % casamino acids (Bacto^TM^, Difco Laboratory^)^, 1 ml of 1M MgSO_4_ (Carl Roth GmbH + Co. KG), 3 g agar (Agar-agar, Kobe I, Carl Roth), 200 ml of 5X M8 solution (64 g Na_2_HPO_4_.7H_2_O (Carl Roth GmbH + Co. KG), 15 g KH_2_PO_4_ (Carl Roth GmbH + Co. KG), 2.5 g NaCl (Acros Organics)) in 90-mm Petri plates which were incubated in upright position for four to five days at room temperature. The experiment was replicated independently four times for R47 and three times for R32. For exposure experiments, split Petri dishes were used (Figure S7). LB agar was poured in one compartment of the dish and 0.3% M9 minimal agar medium was poured in the other compartment. Ten ml of medium was poured in each compartment of the Petri dish. The LB agar side was inoculated with three 10-µl drops of OD adjusted bacterial suspension while the M9 agar side was pricked at the center with a pipette tip. The LB side of the plate was inoculated 6h before inoculation on the M9 agar side. Plates were sealed with parafilm and incubated at room temperature for two days. Pictures were taken and analyzed using the ImageJ software (version 1.53i) (40) to obtain the growth area in pixels square.

### Biofilm formation assays

Bacterial collection and inoculation in KB broth were as described in ‘*Growth curves and pyoverdine measurements*’. The inoculated plates (96-well plates, Corning, NY, USA) were statically incubated at 30 °C for 48h and biofilm measurements were done as previously reported (41) using a Cytation5 plate reader (BioTek, Winooski, VT, USA). The experiment was replicated three times in total with 36 technical replicates each. For exposure experiments, 24-well plates (Corning, NY, USA) were used. 100 µl of OD adjusted bacterial cells were added to 1 ml of LB and LB plates were sealed with parafilm and incubated for 48h statically at 30 °C. A schematic of the experimental setup is shown in Figure S8. Exposure experiments were independently reproduced three times with 6-20 technical replicates.

### RNA extraction

OD adjusted bacterial cell suspension were used to inoculate 6-well plates (Corning, NY, USA) at a final OD600 of 0.1 in KB broth. The plates were then sealed with parafilm. The experimental setup is shown in Figure 3a. For all three time points, the culture volume needed to obtain cells equivalent to OD600 = 3 was collected from the middle two wells (dark boundary in Figure 3a) by centrifugation at 5000 rpm for 5min at 4 °C. Samples from each treatment and time point were collected from three independent experiments. The collected cells were flash frozen using liquid nitrogen and stored at −80 °C until RNA extraction. RNA extraction was performed using standard phenol-chloroform extraction with the Trizol method (42). Samples were treated with DNaseI (Sigma Aldrich) and then re-extracted using the same Trizol method as above. The RNA was quantified and analyzed by the NGS facility in Bern (using both Qubit and Bioanalyzer quantification). The libraries were prepared according to the TruSeq Stranded mRNA Sample Preparation Guide (Illumina). Briefly, total RNA was purified, fragmented, reverse transcribed, and amplified to generate the libraries, which were subjected to high-throughput 100bp single-end sequencing on a NovaSeq 6000 instrument (Illumina).

### RNA-seq data analysis

Quality control of the reads was performed with FastQC v0.11.7 (fastqc: https://www.bioinformatics.babraham.ac.uk/projects/fastqc/), revealing excellent quality reads for all samples. The reads were cleaned using fastp v0.19.5 (43) with a special focus on removing polyG trails and keeping only full length reads (100bp). The reference genome for *Pseudomonas chlororaphis* R47 (CP019399) was downloaded from the NCBI data analysis and indexed for STAR v2.5.0b (44). Then the reads of each sample from step 1 were remapped to genes with STAR using the annotation information (gtf file). The final table of counts was obtained by merging the individual tables with unix commands. The read counts were then analyzed using the R library DESeq2, version 1.30.1 (45). The Log2 (fold change) cut-off was set to greater than or equal to 1 and the adjusted p-value threshold was set to less or equal to 0.05.

### qPCR analysis

Three biologically independent samples from R47 wild type strain collected for RNA-seq were analysed for the expression of the *hcnA* gene. Primers used are described in Table S3. RNA samples were reverse transcribed using the SensiFAST cDNA synthesis kit (Bioline). Quantitative PCR was performed according to the manufacturer’s instructions (Bioline). Briefly, 25 ng of cDNA was mixed with 0.5 µl of forward and reverse primers (10 µM), 1.5 µl of sterile water and 7.5 µl SYBR Hi-ROX mix (Bioline). Reaction conditions were as follows; 95 °C for 15min (initial denaturation), 45 cycles of 95 °C for 15s, 60 °C for 15s, 72 °C for 30s. Data were analysed using the delta Cq method.

### Volatilome analysis by gas chromatography coupled to mass spectrometry

Volatile compounds emitted by wild types and mutants of the strains R32 and R47 were collected as described in (46). Briefly, three single bacterial colonies were used to inoculate LB and cultures were grown at 30 °C overnight. On the following day, the cultures were adjusted to OD600 = 1 and 100 µl were spread onto 5-cm glass plates with 5 ml prepoured LB agar. The plates were incubated at room temperature for 16h. The plates were then shifted to volatile collection chambers and volatile compounds were collected with a charcoal filter for 48h as described in (37). The collected volatiles were then extracted from the charcoal filter using 75 µl of dichloromethane (VWR) and stored at −80 °C until further analysis with GC/MS using the parameters described in (46). The acquired chromatograms were then used for extracting mass features (mz@RT) using MZmine-2.20 (47). The extracted data was then used for statistical analysis using Metaboanalyist (48). The data was normalized with log-transformation and automatic scaling. To identify statistically different mass features between wild type and HCN-deficient strains, student’s *t*-test was used with a P-value cutoff set at 0.05. The mass features detected in uninoculated LB agar were removed before statistical analysis. The experiment was replicated three times independently.

### Hydrogen cyanide quantification

R47 and R32 wild type strains were inoculated and grown in 6-well plates as described in the *RNA extraction* section except that 1 ml 0.38 M NaOH (Merck, Germany) was added to the middle two wells to capture HCN in the surrounding. At the transition phase between the exponential and stationary growth phases, the experiment was stopped, the OD600 was measured using the Cytation5 automated plate reader, and the NaOH samples were collected. In a 96-well plate, aliquots of 100 µl of collected NaOH samples were mixed with 100 µl of 1 M HCl (Fisher Scientific U.K. Ltd) and immediately covered with HCN detection filter paper (16). After 30 min incubation at room temperature, the filter paper was scanned using a Epson Perfection V370 photo scanner and analysed using ImageJ (version 1.53i). A standard curve was made with KCN (Merck, Germany) solutions at 31, 62 and 125 µM in 0.38 M NaOH. The experiment was replicated three times independently.

### Statistical analysis

All statistical analyses for phenotypic assays were performed using Graphpad Prism 8.0.1 (https://www.graphpad.com/). For pair-wise comparisons, the Mann-Whitney test was done (Figure 1c-e). For multiple comparisons, the linear model assumptions were checked and ANOVA followed by Tukey’s test was used for analysis (data was log transformed for Figure 3d).

## Acknowledgements

Financial support from the Swiss National Science Foundation (grants 179310 and 207917 to LW) is gratefully acknowledged. We also thank Mout de Vrieze for her help with statistical analyses. LW and AA conceived the project and wrote the manuscript, with help from CK. AA performed all experiments, with help from FLH. LF performed transcriptomics analysis, EAM performed HPLC analysis. All authors commented on the manuscript. The authors declare no competing interests.

