## Supplemental figures for "Biological hydrogen cyanide emission globally impacts the physiology of both HCN-emitting and HCN-perceiving *Pseudomonas*"

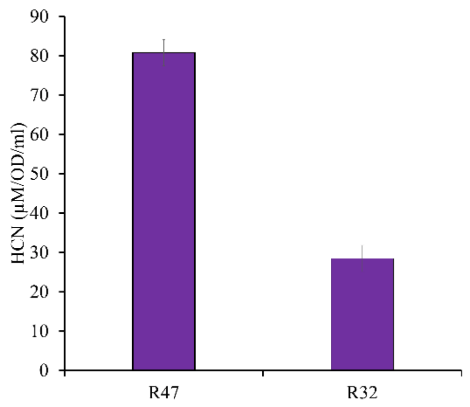


**Figure S1. Hydrogen cyanide quantification.** Wild type strains of *Pseudomonas chlororaphis* R47 and *Pseudomonas putida* R32 were inoculated in 6-well plates (two wells on the left and two on the right while the two centre wells were filled with NaOH to capture HCN) for 7h at 30 ˚C with shaking. After 7h, NaOH was collected and HCN was quantified as described in the experimental procedure section. Bars represent the average HCN concentration in 1 ml bacterial culture ($OD_{600} = 1$) using three independent experiments. Error bars represent standard errors.

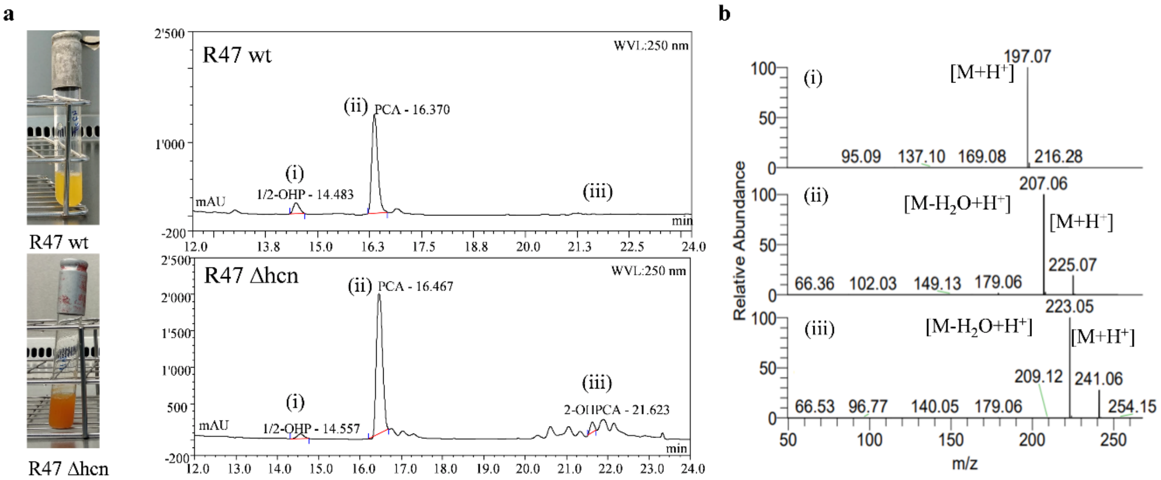


**Figure S2. HCN represses the production of phenazines. a,** Phenazine quantification assay using an HPLC-UV method as described in the experimental methods section. Images represent *Pseudomonas chlororaphis* R47 wt and R47 Δhcn in liquid culture in King's B medium after 24h growth at 30 °C with shaking. Chromatograms represent the HPLC-UV profiles with the x-axis as time in minutes and the y-axis as peak intensity (arbitrary units). **b,** MS profile for peaks (i), (ii) and (iii). Peak (i) = 1 or 2-hydroxyphenazine, peak, (ii) = phenazine-1-carboxylic acid (PCA) and peak (iii) = 2-hydroxyphenazine carboxylic acid.

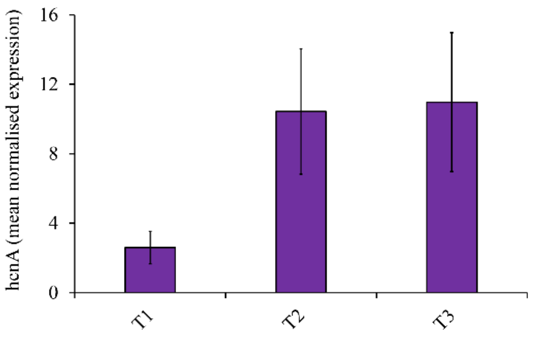


**Figure S3. *hcnA* expression starts at the early exponential growth phase in *Pseudomonas chlororaphis* R47.** qPCR analysis of the expression of the *hcnA* gene from the HCN biosynthetic operon. *rpoD* was used as reference gene. RNA samples and time points used for this analysis were the same as those used for transcriptomic analysis (Figure 2a). Bars represent the mean normalized expression compared to *rpoD* expression. Bars represent the average from three independent experiments for each time point. T1; early log phase, T2; transition phase (exponential to stationary), T3; stationary phase.

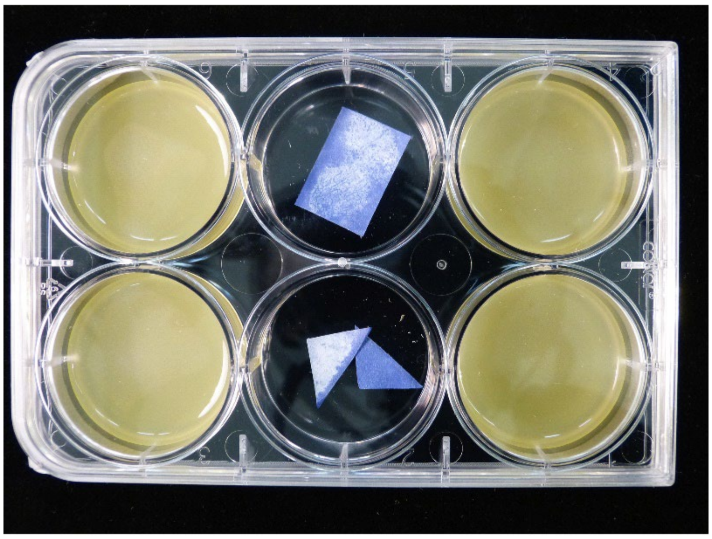

**Figure S4. Volatile-mediated exposure assay in 6-well plates.** The two wells on the left and on the right, respectively, were inoculated with the *Pseudomonas chlororaphis* R47 wild type strain. Filter papers imbibed with copper (II) ethyl acetate and 4,4-methylenebis(dimethylaniline) solution for HCN detection were placed in the two middle wells. The cultures were incubated at 30 ˚C for 18h with shaking before capturing the image. The blue color in the middle two wells reveals the presence of HCN.

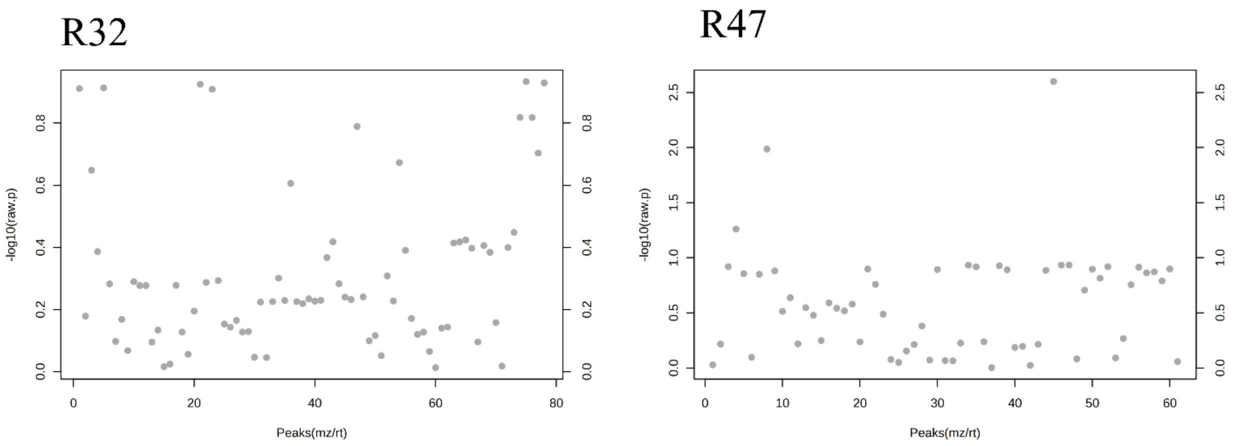


**Figure S5. Volatilome of *Pseudomonas putida* R32 and *Pseudomonas chlororaphis* R47.** The wild type strains and the respective HCN-deficient mutants were grown on LB medium for 48h. The volatile compounds they emitted were collected using a closed-loop stripping method and analysed using GC/MS. Dots represent mass features. Grey color represents mass features that were not statistically different between the wild types and their respective HCN-deficient mutants. A Student's T-test with a P-value cutoff set at 0.05 was used to find statistically different mass features between the wild types and the HCN-deficient mutants. On the x-axis, mass features are plotted against the –log10(raw P value) on the y-axis. Mass features present in LB medium controls were removed. Data represent mass features detected in three independent biological replicates. Note that HCN cannot be detected by standard GC/MS protocol and is therefore not depicted in the graph.

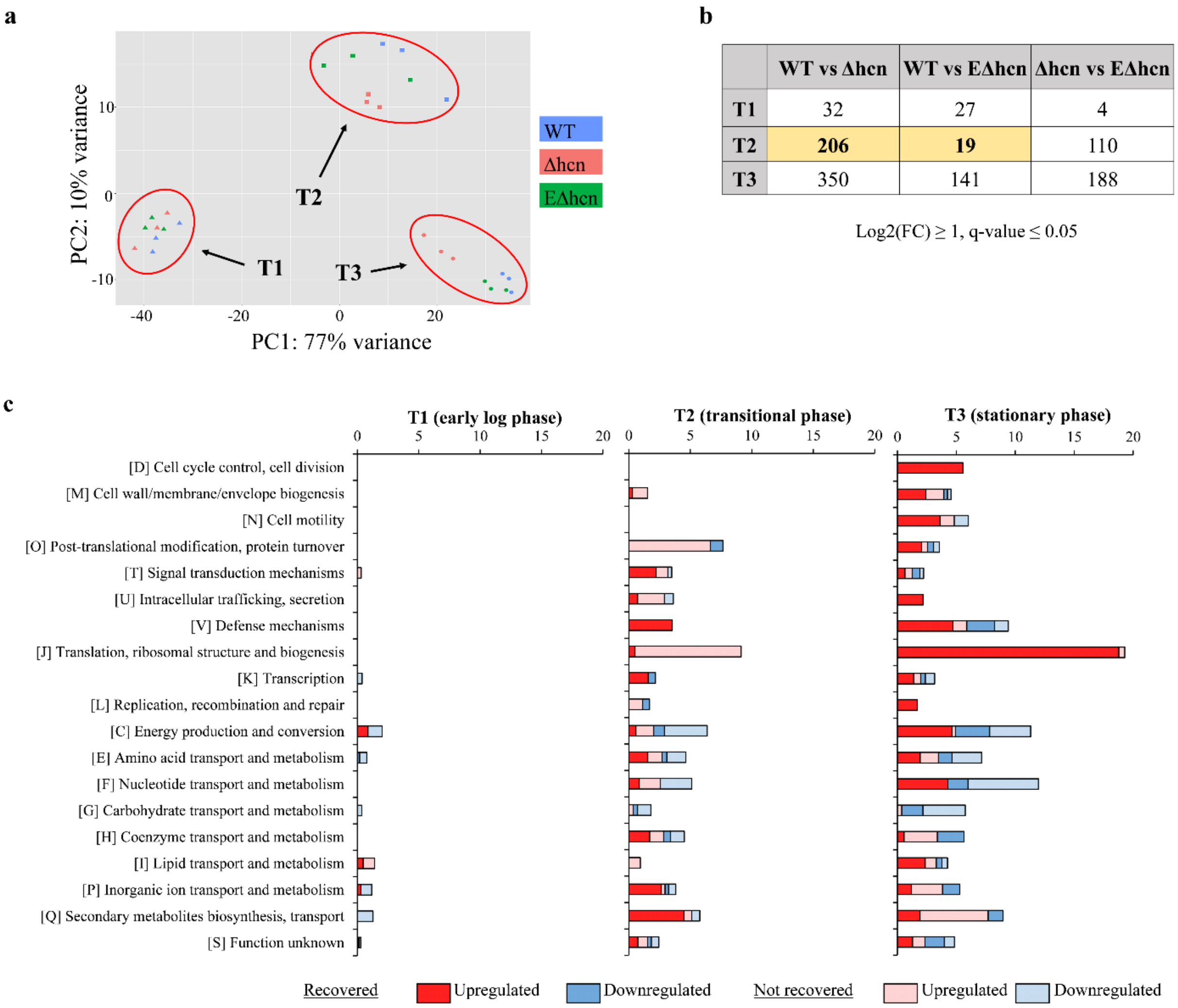


| | WT vs Δhcn | WT vs EΔhcn | Δhcn vs EΔhcn |
|---|---|---|---|
| T1 | 32 | 27 | 4 |
| T2 | **206** | **19** | 110 |
| T3 | 350 | 141 | 188 |

**Figure S6. Exposure to exogenous HCN from *Pseudomonas chlororaphis* R47 wild type leads to global transcriptomic reprogramming in the mutant. a,** Principal component analysis of transcriptomic data of the wild type strain (wt), the hcn mutant strain (Δhcn) and the hcn mutant strain exposed to HCN emitted by the wild type strain (EΔhcn) at time points 1, 2 and 3. T1; early log phase, T2; transition phase (exponential to stationary growth), T3; stationary growth phase. **b,** Dysregulated genes in wt vs Δhcn, wt vs EΔhcn and Δhcn vs EΔhcn comparisons. Log2 (fold change) threshold was set to 1 and q-value to 0.05. **c,** COG category analysis of dysregulated genes in Δhcn compared to wt as shown in Figure 2c, with supplementary data from the comparison between wt and EΔhcn to highlight recovered vs. non-recovered gene expression in EΔhcn. A gene was considered recovered if the log2 (fold change) expression value in 'wt vs EΔhcn' comparison was less than 50% of its value in 'wt vs Δhcn' comparison.

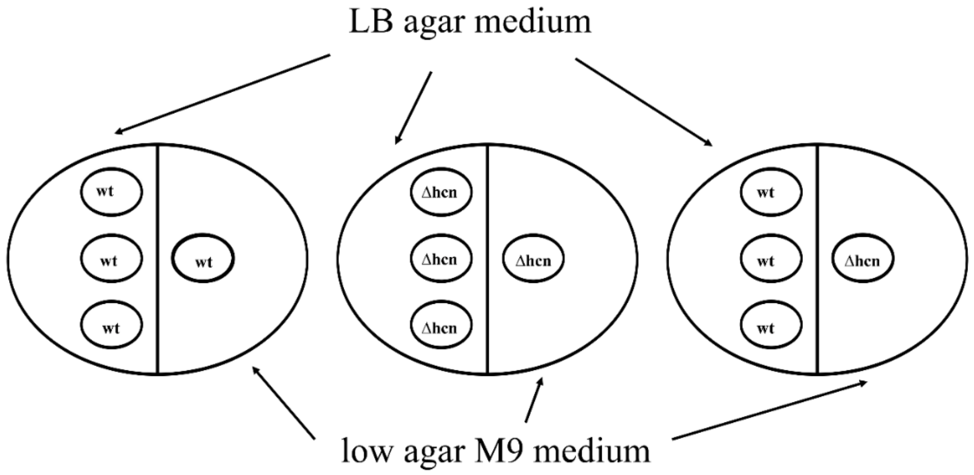


**Figure S7. Split plate exposure experimental setup to measure motility.** The left side of the split petri dish was inoculated with three drops of bacterial suspension 6h before the inoculation on the right side of the plate as described in experimental procedures. The left side was filled with LB agar medium while the right side was filled with low agar medium. The plate on the left represents the wild type strain exposed to the wild type strain, the plate in the middle represents the HCN-deficient mutant exposed to the HCN-deficient mutant, while the plate on the right shows the HCN-deficient mutant exposed to the wild type.

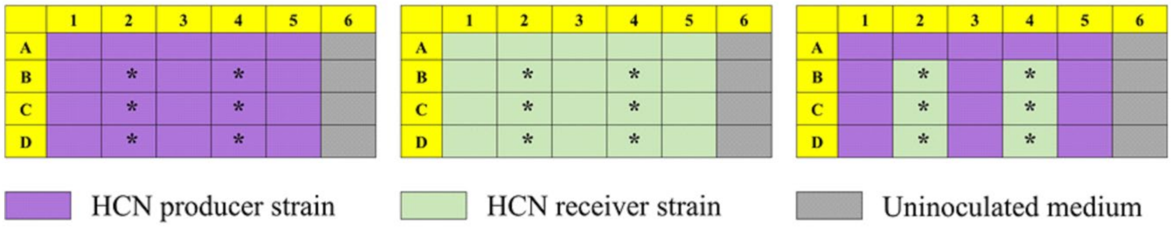


**Figure S8. 24-well exposure experimental setup to measure biofilm formation.** HCN producing bacteria were inoculated in the wells colored in purple, while the HCN receiving bacteria were kept in green wells. Grey wells indicate uninoculated wells. Wells marked by * were sampled for biofilm analysis.
